# THE CEREBELLUM CONVERTS INPUT DATA INTO A HYPER LOW-RESOLUTION GRANULE CELL CODE WITH SPATIAL DIMENSIONS: A HYPOTHESIS

**DOI:** 10.1101/2023.07.14.548987

**Authors:** Mike Gilbert, Anders Rasmussen

## Abstract

We present a theory of the inner layer of the cerebellar cortex, the granular layer, where the main excitatory input to the cerebellum is received. We ask how input signals are converted into an internal code and what form that has. While there is a computational element, and the ideas are quantified with a computer simulation, the approach is primarily evidence-led and aimed at experimenters rather than the computational community. Network models are often simplified to provide a noiseless medium for sophisticated computations. We propose, with evidence, the reverse: physiology is highly adapted to provide a noiseless medium for straightforward computations. We find that input data are converted to a hyper low-resolution internal code. Information is coded in the joint activity of large cell groups and therefore has minimum spatial dimensions – the dimensions of a code group. The conversion exploits statistical effects of random sampling. Code group dimensions are an effect of topography, cell morphologies and granular layer architecture. The activity of a code group is the smallest unit of information but not the smallest unit of code – the same information is coded in any random sample of signals. Code in this form is unexpectedly wasteful – there is a huge sacrifice of resolution – but may be a solution to fundamental problems involved in the biological representation of information.

## 1. Introduction

It is generally assumed that brain-coded information is rich and replicable with computer-like high fidelity, and that those features make it precise and allow computations to be powerful. In any event, this is not regarded as contentious. In many neural network learning models, including cerebellar models, computations are implemented by learned synaptic changes that selectively adjust synaptic transmission strength. This has been a mainstay of much modelling for 50 years. We will argue, *inter alia*, that these influential ideas may need to be reconsidered.

The inner layer of the cerebellar cortex, the granular layer, receives the main excitatory input to the cerebellum, from mossy fibres. Mossy fibres contact both granule cells and large inhibitory interneurons, Golgi cells. Golgi cells in turn inhibit granule cells. The granule cell axon rises into the outer layer, the molecular layer, where it divides in two to form parallel fibres, which lie parallel to the cerebellar surface and other parallel fibres. Parallel fibres make contact in passing on Purkinje cells, which carry the sole output of the cerebellar cortex.

We attempt to explain neuroanatomy of the granular layer as a strategy that automates the granular layer computation. The computation, in this contention, is the passive result of anatomy (cell morphologies and neuroanatomical architecture) and linear signalling. Speaking generally, neuron-to-neuron transmission involves the complex interaction of a large array of often co-dependent biophysical and other variables which are specific to the cell types making and receiving contact. Yet despite the potential to be dysfunctionally noisy, linear relationships are in fact conserved. There is now a substantial body of evidence that rate-coded information is linearly conserved in communication between cerebellar neurons [1–7], and that cells are adapted for faithful transmission [8–14]. Firing rates that linearly code behavioural and task metrics have been observed ‘at all levels of the cerebellar circuit’ [15 p.239, citing 30 sources]. There are a number of examples that the cerebellum is adapted to disengage or otherwise compensate for intrinsic non-linear properties of neurons and synapses [12, 13, 16, 17]. This is – we submit – part of a general strategy of isolating the effect of parameters that code information while eliminating or mitigating interference by other variables. Evidence of linear signalling in the granular layer is particularly rich. We include a short review in a dedicated section before the Discussion to which we cross refer from the main text.

We will argue that the granular layer is in effect the physiological form of a network of linear units which randomly sample firing rates in the afferent cell layer, within topographically-defined boundaries. As sampling units have identified anatomical counterparts, anatomy provides model parameters. We field evidence in the main text to explain the derivation of model parameters. Parameter values (usually a range) are usually available from the literature.

Classically, neural network models solve a problem. The nature of the problem – and therefore the function of the network – is a proposal and becomes an assumption of the model. The solution is a computational mechanism; the mechanism is the model. The direction is function → mechanism (the first two steps of Marr’s three levels of analysis; the third step is to consult the evidence for support). We work in reverse. We propose first a mechanism that can explain the evidence, then infer function from the mechanism. In our process, high-level ideas come last. We use modelling in a different role, to quantify and test the ideas by simulating the mechanism. In our approach, the hypothesis and simulation are both the mechanism/model. The simulation is the hypothesis in quantified form. While there is a computational element, and the ideas are quantified with a computer simulation, it is not primarily a computational paper.

Note that we do not propose to be mathematically interesting. The heart of the proposal is that the physiological computation is in fact not sophisticated, against expectation – at least, there is a plausible argument that this is the case, which we present as an alternative to current models. For that reason, physiology has a high profile in the paper. The sophistication of the biological design is found in the fact that physiology is adapted to isolate computationally relevant parameters (and computations themselves) from interference by other variables. Biophysical events which accomplish this result are complex. If we are correct, a future challenge will be to add them, so that the model is populated with a smaller scale of detail. Here, however, they are superfluous.

The next section describes our methodology in more detail. We encourage the reader to read the first subsection (2.1 ‘The relationship of physiology and model parameters’) before proceeding because it describes how the computation is related to the physical hardware of the system.

## 2. Methods

### 2.1 The relationship of physiology and model parameters

#### 2.1.1 SUMMARY OF METHODOLOGY

We propose a physiologically detailed hypothesis which we recreate in silico in the form of a network of unit functions. Functions are attributed to identified anatomical hardware, connected by known anatomy, and the functions themselves are individually evidenced. In this form, we are able to model large cell networks in high *relevant* definition, without simplifying assumptions. ‘Relevant’ is italicised to emphasise that parameter values represent data – numbers used to code information – and not biophysics. Hence the claim that the *computation* is not simplified.

We argue that the computation is a passive effect of anatomy. The simulation provides a demonstration that processing by anatomy provides a mechanism that (i) self-regulates the fraction of granule cells that fire at any time, and (ii) converts input signals into an unexpectedly low-resolution internal code with spatial dimensions.

#### 2.1.2 COMPUTATIONAL ARCHITECTURE

The granular layer receives input to the cerebellum and converts it to an internal code.

Input is received from mossy fibres. Mossy fibres contact and excite granule cells and Golgi cells. Golgi cells inhibit granule cells. Excitation of Golgi cells and excitation and inhibition of granule cells all happen in a structure called a glomerulus.

Granular layer anatomy is the physical form of an orderly network of computational units. A unit takes inputs and returns output. Inputs and output represent values (that is, numbers). A unit is the site of biophysical events which relate inputs and output – the unit function.

Physically, the granular layer appears anatomically seamless – a carpet of cells. But they are wired so that units are organised functionally in layers. The outputs of a layer provide the data sampled (i.e., received as inputs) by units in the next layer.

Data are constantly refreshed. Computations in a layer run in parallel, continuously, so unit outputs are analogue. This occurs in all layers at once. The aggregate of computations relates input and output of the system: the (granular layer) network computation.

All layers of unit functions span the network, forming a long, thin strip. All strips are sagittal, and therefore parallel. Information is coded collectively in the concurrent outputs of a layer. The code is a frequency distribution, so that a whole layer, at any given instant in time, codes information that is both indivisible (cannot be subdivided into smaller units of information) and yet coded in any random sample of unit output values. The biophysical form of outputs depends on the layer.

#### 2.1.3 HOW ARE UNITS CONNECTED?

By anatomy, so we know what the connections are, because anatomy has been reported in detail. The number of inputs to a unit is usually anatomically randomly variable in a reported range. In some cases, we simulate that by generating the number with a distributed probability, and in others we use the mean.

It is unnecessary to know which individual cells contact each other because (biologically and in the model) contact is at random inside topographically defined dimensions. This has the asset for modelling that a target population can be represented by randomly sampling data received as input to a location.

Biologically, a location is a volume with spatial dimensions. *In silico*, we can represent that as a population of unit functions. We know from anatomy (or can derive) the size of unit populations, population ratios, and convergence and divergence ratios.

#### 2.1.4 WHAT DETERMINES THE SHAPE AND SIZE OF A POPULATION OF UNIT FUNCTIONS?

The cerebellar cortex is anatomically seamless — there are no physical boundaries that divide cells into groups, just organisation that reflects topography. Microzones are an example (microzones are defined by their climbing fibre input) [18].

Microzones are divisions of the molecular layer. Mapping studies suggest that the granular layer may have similar organisation, but the picture is less clear [19]. However, the morphology and termination pattern of mossy fibres suggest a functional division into strips that may mirror microzones.

Mossy fibres give rise to sagittally-extending collaterals, the direction of the long axis of microzones (Fig 1). Collaterals branch terminally, and each branch ends in a cluster of terminals, so that a single cell terminates in a spaced-out row of terminal clusters which are always lined up in the same direction (parallel to microzones but in the granular layer) [20–23]. Clusters are separated by a randomly variable distance typically not less than 200 µm [20, 21] measured from edge to edge of clusters (so they are separated by a greater distance measured from centre to centre). Terminals are small compared to the average volume occupied by a cluster, and intimately and randomly intermingled with terminals of other mossy fibres.

**Fig 1.**
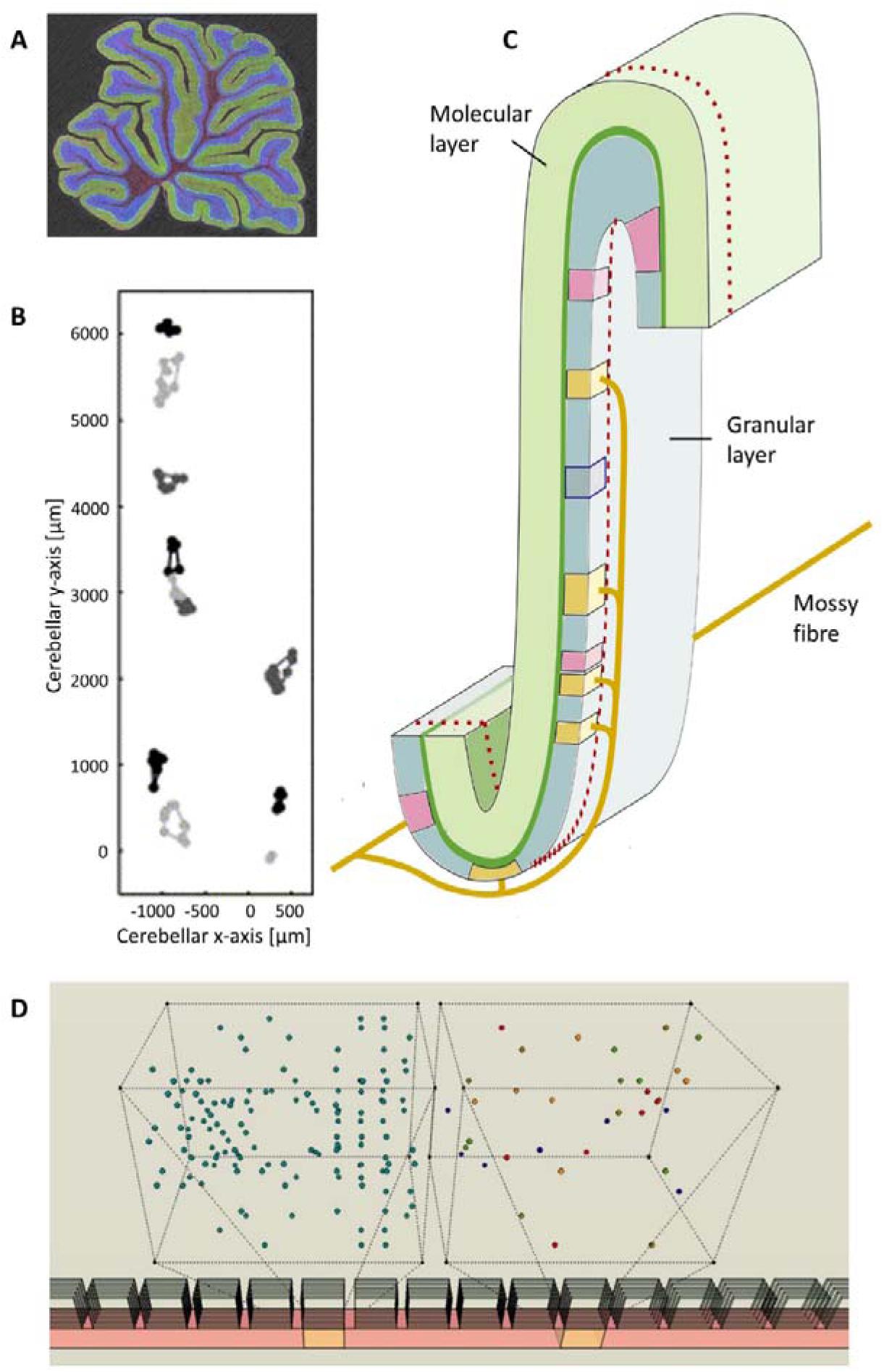
Mossy fibre termination. **A** Sagittal section of the rat cerebellum (obtained by multi-photon microscopy, courtesy Tom Deerinck). The outer layer of the cerebellar cortex, the molecular layer, is stained green and the inner layer, the granular layer, is blue. **B** Output of a procedure used to group mossy fibre terminals into clusters [21 fig 6], reproduced with permission. All terminals belong to one mossy fibre, which terminates in two strips. Clusters are shown in different shades of grey. Cerebellar x-axis: parallel fibre axis; Cerebellar y-axis: parasagittal axis. **C** Schematic of an area of the folded cerebellar cortex. The strip inside the red dotted line is the width and depth of a microzone but only part of its length. Yellow zones are terminal locations of a single mossy fibre. Pink zones are terminal locations of a second mossy fibre. In reality, zones overlap extensively and terminals are intermingled. The zone enclosed by a blue line has the dimensions of a ‘field’, a nominal division of a rank with the average dimensions of a mossy fibre terminal cluster. **D** Section of a microzone viewed obliquely from the cerebellar surface, to illustrate the size and random intermingling of mossy fibre terminals. The wire frame boxes are an exploded view of the orange fields. The right box shows randomly distributed terminals of 5 mossy fibres (each a different colour). Proportions of terminal size to field size are preserved. The left box shows terminals of 20 mossy fibres (out of ∼180 that contribute terminals to a field).

Accordingly, mossy fibres do not terminate at a point but a region with dimensions: the field occupied by a terminal cluster. A single mossy fibre gives rise to several terminal branches and each of those to several terminals, such that a sagittal strip of the granular layer receives multiple copies of every signal at each of multiple locations.

The dimensions which enclose a cluster (cluster size) are variable (because it depends on the number of terminals in a cluster), averaging 200 µm sagittally x 150 µm mediolaterally [21]. Say we divide the granular layer into strips and strips into ‘fields’ (each perhaps 1/100^th^ part of a strip, but still each containing thousands of granule cells). Fields are average cluster size. Fields are simply nominal, a convenient modelling device. But strips are functional – the minimum dimensions at which the cerebellum makes any effort to tell signals apart, we submit.

The shape and size of a population of unit functions are a consequence of this topography – the dimensions of a strip where mossy fibre terminals are mixed up at random. This is the first way that mossy fibre morphology is the key to the cerebellar code. A sagittal row of 75–100 fields has the area (viewed from the cerebellar surface) of a mid-range microzone [18, 24–26]. We term a row of fields a ‘rank’ (strip has been used in the literature to refer to other divisions of the cerebellar cortex).

#### 2.1.5 SAMPLING WITH REPLACEMENT

##### 2.1.5.1 Terminal branching mimics independent sampling by a field of input rates to a rank

We propose that terminal branching of mossy fibre collaterals simulates independent random sampling, by fields, of input rates to a rank. That is, inclusion of a rate in a sample does not affect the probability that it is including in any other sample, also called sampling with ‘replacement’.

If each mossy fibre could terminate in only one field, a mossy fibre ‘sampled’ by one field clearly cannot also be sampled by any others. Termination by a single mossy fibre in a random number of terminal branches, at multiple locations, removes that restriction.

In theory, sampling with replacement would mean any number of fields can receive the same signal, up to all of them. Clearly this is not true – in reality there is an anatomical limit. However, even if there was real replacement, the number of repeat samplings is constrained by a probability distribution – effectively, a cap. If, then, the number of terminal branches per mossy fibre has the same probability distribution, branching has the same result as if firing rates received by each field in a rank were in fact a random sample with replacement of firing rates received by the rank as a whole.

We can test that by calculating the probability distribution in the hypothetical case that mossy fibres rates were in fact sampled with replacement but with a probability (that a given sample contains a given rate) that reflects the average number of terminal branches per cell. The probability, *p*, that a given rate is selected *k* times in taking *N* samples, sample size *x,* assuming mossy fibres each give rise to an average of *m* terminal branches, is given by

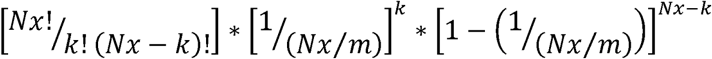

where *N* is the number of fields per rank, each receiving innervation by a number of mossy fibres, *x*, given by the derivation of mossy fibre rates in section 2.2, and assuming *m* = 5. The results are shown in Table 1.

**Table 1.** Probability *p* of *k* repeat selections of a single mossy fibre.

| $k$ | 0 | 1 | 2 | 3 | 4 | 5 | 6 | 7 | 8 | 9 | 10 | 11 |
| --- | --- | --- | --- | --- | --- | --- | --- | --- | --- | --- | --- | --- |
| $p$ | 0.007 | 0.034 | 0.084 | 0.140 | 0.175 | 0.175 | 0.146 | 0.104 | 0.065 | 0.036 | 0.018 | 0.008 |

Although data are scarce, the value of *m* is consistent with what we know about the biological range of the number of terminal clusters per mossy fibre [21, 27]. However, some different value would still make the same point. In hypothetical conditions in which it is theoretically possible for a mossy fibre signal to be selected any number of times, the number is restricted statistically to a small range. The number has the same distributed probability as the number of mossy fibre terminal branches per cell. Physiology in this way mimics sampling with replacement.

##### 2.1.5.2 Terminal clustering mimics independent sampling by single cells of input rates to a field

Granule cells and Golgi cells both receive contact from a random sample of mossy fibres which innervate a field. We propose that mossy fibre terminal clustering simulates independent random sampling, by individual cells, of input rates to a field. The principle is the same here as with terminal branching, but at local, field scale. Mossy fibre terminal clustering, and random intermingling with terminals of other mossy fibres (Fig 1), means that contact on any particular cell does not mean other cells cannot also receive the same signal. Again, to approximate independence, it is only necessary for the number of repeat selections of a signal to have the same probability distribution as if there was actual replacement.

A practical problem with the strategy of terminal clustering is that a target could receive the same signal twice (or more). The risk is mitigated by taking small samples. A sample size of 4 is small enough for a low risk. Granule cells average 4 dendrites, each receiving contact from a single mossy fibre terminal, and we estimate that Golgi cells receive an average of 4 inputs to each of their basal dendrites (see derivation of parameters in later sections). Given *m* x 100 terminals and an independent and equal chance that contact is by any terminal, the probability of an *n* contact by the same mossy fibre on a single dendrite is:

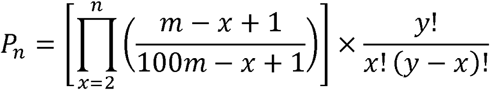

where *y* is the number of inputs to a single dendrite and *m* is the number of terminals per cluster, provided by (for illustration) 100 mossy fibres. Calculated in this way, around one in every twenty samples contains a duplicate, and less than two in a thousand contain more than one. The actual fractions are smaller because the number of mossy fibres that innervate a field is higher, around 180 (see ‘Derivation of mossy fibre rates’ later in this section).

##### 2.1.5.3 The combined effect of terminal branching and terminal clustering

The combined effect is a biological facsimile of simultaneous random sampling with replacement by individual granule cells and Golgi cells of firing rates received as input to a whole rank, notwithstanding that they each receive fixed contact from a tiny fraction of mossy fibres. Note that it is the population of mossy fibre rates that are hypothetically sampled with replacement, and not *signals*. The population of signals includes copies of rates because of branching and clustering.

#### 2.1.6 WHAT IS THE COMPUTATION?

So: the effect of duplicating mossy fibre signals – by terminal branching and clustering – is to mimic sampling with replacement. Granule cells and Golgi cells each individually, effectively (i.e., for granular layer computational purposes), independently randomly sample mossy fibre input rates to a whole rank. This is the second way that mossy fibre morphology is the key to the cerebellar code.

This has the result that unit functions provide the physiological instrument of (continuously) computing (a value linearly related to) the sum of independent random samples of the afferent data layer. Doing this many times (in parallel) generates a distribution with a narrower range and a linearly related mean, by the central limit theorem.

Repeating that layer on layer progressively narrows the data range, while conserving a linear relationship of all the means. The range of granule cell rates is further narrowed because granule cells which fire – a small fraction – are confined (by the glomerular competition – see section 5) to those which receive mossy fibre signals at the higher end of the mossy fibre range.

### 2.2 Derivation of mossy fibre rates

#### 2.2.1 RATIONALE

We run the computation *in silico*. We represent the granular layer as a network of unit computations organised in layers, as described. Support for the size of unit populations, population ratios, and convergence and divergence ratios is cited in context, in the relevant section of the main text. Support for unit functions is discussed in the section before the Discussion.

The simulation is not an additional step in ideas or detail. It is a quantified form of the hypothesis. The hypothesis describes a computational mechanism; the simulation performs the computations. The purpose of the simulation is to confirm that the mechanism is feasible, with plausible evidence. This section describes the derivation of mossy fibre rates received by a rank and by fields taking account of terminal branching and terminal clustering, the randomly variable number of terminals per cluster and random variation of the number of terminals in a cluster which fall inside and outside a field when a cluster straddles a field boundary.

The frequency distribution of mossy fibre rates received as input to a granular layer rank is unreported. We used a large bandwidth in the physiological range and varied the shape of the distribution to confirm that the operation of the mechanism does not depend on the input to it. The shape of the mossy fibre distribution is included in all figures.

#### 2.2.2 THE NUMBER OF MOSSY FIBRES THAT INNERVATE A RANK

Each field receives a random sample mossy fibre firing rates received as input to a rank. To simulate that, we need the size of the sampled population – the number of mossy fibre signals received by a rank – and sample size, the number received by a field.

Despite the repeating ‘crystalline’ architecture of the cerebellar cortex, at local level there are anatomical random variables. Variables are: i) the number of mossy fibres that innervate a field, ii) the number of terminals they each end in, and iii) the number of those that are inside field limits.

The number of mossy fibres afferent to a field has been estimated at 100 [21]. However, that number is a field-sized fraction of the general population. The actual number is larger because mossy fibres end in a dispersed group (cluster) of terminals, and most groups straddle field limits.

We work from an estimate of the total number of terminals per field, which is, in turn, derived from convergent estimates of the number of granule cells per field. (Estimate 1) There are an estimated 1.92 x 10^6^ granule cells per μl [28]. The number of fields which fit in that volume is (1,000/150)^2^ x (1,000/200) = ∼222, assuming a field depth of 150 μm [21]. The number of granule cells in a field is, therefore 1,920,000/222 = 8,649. (Estimate 2) An estimated 350,000 parallel fibres pass through a single Purkinje cell’s territory [28]. Parallel fibres run about 3 mm in each direction from the granule cell soma [29, 30]. The Purkinje cell dendritic arbour fills the molecular layer vertically and averages ∼200 µm wide. Therefore, 6,000/150 = 40 fields fit in a row that provides parallel fibres that pass through a Purkinje cell dendritic territory, giving 350,000/40 = 8,750 granule cells per field. Estimates vary of the number of granule cells which extend a dendrite into a single glomerulus. 50 is mid-range. (8,750/50) x 4 = 700 glomeruli per field.

Granule cells have 3–5 dendrites, averaging 4, and receive input to each of them from a single mossy fibre (so an average granule cell receives 4 inputs). Estimates of divergence of a single mossy fibre terminal onto granule cells vary [1]. It is unclear how much this reflects biological diversity versus differences in opinion and methods. Here we use a mid-range estimate of 1:50 [31, 32]. Assuming a mossy fibre terminal receives a single granule cell dendrite from each of d granule cells, and given an average of 4 dendrites per granule cell and z granule cells per field, the total number of mossy fibre terminals in a field is *N* = (*z* x 4)/*d* = 700 given *z* = 8,750 and *d* = 50. A convergent estimate is provided by multiplying the per field number of mossy fibres by the average number of terminals per cluster.

Cluster size varies. We take the range as 4–12 terminals per cluster [21]. We assume the number varies at random and cluster sizes occur with equal probability. A maximum of 7 fits in a field (because a field shares the dimensions of the area that encloses a mean cluster). Fields are not anatomically bounded, so many clusters straddle field limits. To reflect that, we assume there is a 1/7 probability that a 7-terminal cluster (or larger) has 7 terminals inside a field and also the same probability of any other number. A 6-terminal cluster has a 2/7 chance that all 6 terminals are inside and a 1/7 chance of any other number, and so on.

Table 2 shows the derivation of the expected (i.e., predicted by probability) ratios of terminals contributed to a field by each cluster size (R1), not counting terminals that lie outside when a cluster straddles field boundaries; and the ratios of mossy fibres contributing *n* = 1–7 terminals (R2), regardless of cluster size. R1 ratios are given by the sum of numbers in each row. So, for example, the ratio of cells that end in 8 terminals to cells that end in 5 is 28:25. R2 ratios are given by the number of appearances of each number in the grid – so the expected ratio of cells which contribute 7 terminals to the number which contribute 4 is 6:12. The ratios effectively give the distributed probability of possible states. For example, there is a probability of 10/(6 + 8 + 10 + 12 + (3 x 9)) = 0.159 that a mossy fibre contributes 5 terminals to a field (regardless of cluster size). Similarly, there is a probability of ((6 x 28)/((6 x 28) + 27 + 25 + 22))/7 = 0.099 that a given cell contributes 7 terminals to a field.

**Table 2.** Derivation of expected ratios.

| size | number of terminals inside |  |  |  |  |  |  | R1 |
| --- | --- | --- | --- | --- | --- | --- | --- | --- |
| **12** | 7 | 6 | 5 | 4 | 3 | 2 | 1 | **28** |
| **11** | 7 | 6 | 5 | 4 | 3 | 2 | 1 | **28** |
| **10** | 7 | 6 | 5 | 4 | 3 | 2 | 1 | **28** |
| **9** | 7 | 6 | 5 | 4 | 3 | 2 | 1 | **28** |
| **8** | 7 | 6 | 5 | 4 | 3 | 2 | 1 | **28** |
| **7** | 7 | 6 | 5 | 4 | 3 | 2 | 1 | **28** |
| **6** | 6 | 6 | 5 | 4 | 3 | 2 | 1 | **27** |
| **5** | 5 | 5 | 5 | 4 | 3 | 2 | 1 | **25** |
| **4** | 4 | 4 | 4 | 4 | 3 | 2 | 1 | **22** |
| **R2** | 6 | 8 | 10 | 12 | 9 | 9 | 9 |  |

If R1 ratios are denoted by a series *x*_4_, *x*_5, …_*x*_12_ and a given element in the series is *x_j_*, the fraction of terminals provided by a given cluster size is *x_j_*/*(x*_4_ *+ x*_5_ *+ x*_12_), and this fraction is multiplied by *N* to give the number of terminals. The number of terminals is divided by the expected mean number of terminals that fall inside field limits (for that cluster size) to give the expected number of mossy fibres that provide them. The number of *j* terminal mossy fibres that innervate a field is thus given by

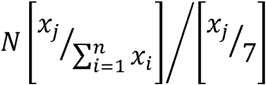

The results are shown in Table 3. The equal number of mossy fibres (mfs) in each column is what we would expect, because we made it an assumption that all sizes occur with equal probability, and mf values are expected values – i.e., in proportions predicted by their probability. The mf total is the expected number of mossy fibres afferent to a field (which contribute at least one terminal).

**Table 3.**
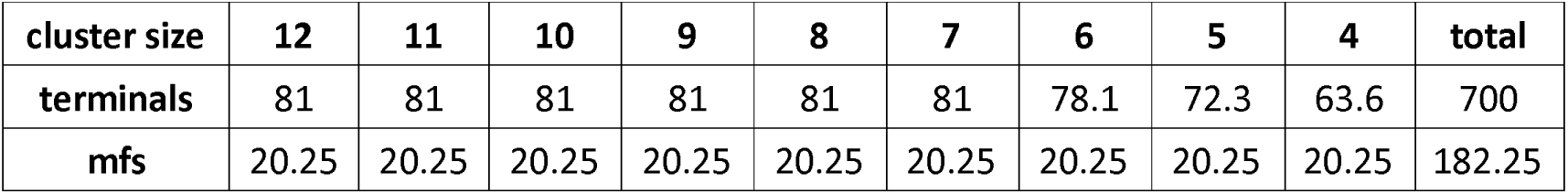
The expected number of mossy fibres which innervate a field.

R2 ratios – the relative incidence of afferent mossy fibres that contribute 7 terminals to a field, 6 terminals, 5 and so on, down to 1 – and the expected number of mossy fibres that contribute them, are shown in Table 4.

**Table 4.** The number of mossy fibres which contribute *n* = 1–7 terminals to a field.

| n | 7 | 6 | 5 | 4 | 3 | 2 | 1 | total |
| --- | --- | --- | --- | --- | --- | --- | --- | --- |
| R2 | 6 | 8 | 10 | 12 | 9 | 9 | 9 | 63 |
| R3 | 42 | 48 | 50 | 48 | 27 | 18 | 9 |  |
| mfs | 17.34 | 23.15 | 28.92 | 34.72 | 26.04 | 26.04 | 26.04 | 182.25 |
| %age | 9.52 | 12.69 | 15.87 | 19.05 | 14.29 | 14.29 | 14.29 |  |

R3 is *n* x R2, giving the ratio of terminals provided by mossy fibres with n terminals inside a field. %age is given by (R2/63) x 100. this is the percentage of mossy fibres with *n* terminals inside a field.

From here, the number of mossy fibres that innervate a rank is given by 182.25 x 100 divided by the mean number of clusters per mossy fibre per rank. If a rank receives, on average, five clusters per mossy fibre, ∼3,645 mossy fibres innervate a rank. This is the estimate used in the simulation unless stated otherwise.

*Assumptions.* Cluster sizes occur with equal probability. The probabilities of each outcome when a cluster straddles field boundaries (how many terminals are inside and how many outside) are as stated. A single mossy fibre ends in an average of 5 terminal clusters per rank.

#### 2.2.3 MOSSY FIBRE INPUT RATES TO A FIELD

Terminal clustering means a field receives multiple copies of most signals. The proportions of mossy fibres afferent to a simulated field that contribute each possible number of terminals (1–7) are generated in simulated steps. In the first step, the simulation randomly generates, for each field, the cluster size ratio. This means, the relative proportions of afferent mossy fibres (which contribute at least 1 terminal to a field) that end in 4 terminals, 5, 6 and so on up to 12, assuming they occur with equal probability, disregarding how many terminals fall inside and outside field boundaries. In step two, the ratio of cluster sizes is weighted to reflect the greater probability that large clusters contribute more terminals to a field. In step 3, ratios are converted to the fraction of terminals contributed by each cluster size. When multiplied by the total number of terminals, this gives the number of terminals contributed by each cluster size. In step four, the values obtained in step three are divided up between cells that contribute a single terminal, 2 terminals, 3 and so on, to reflect straddling of field limits. The proportions give the number of mossy fibres afferent to a simulated field that contribute each number of terminals up to seven, the maximum.

There is no field-level effect of mossy fibre terminal branching. Terminal clusters arising from the same mossy fibre are separated by a minimum physiological distance, so that a field cannot contain terminals of more than one cluster per mossy fibre.

Fields receive a randomly variable number of copies of most signals in the cerebellum and the simulation. In the simulation, copies are added after sampling of mossy fibre rates to obtain rates received by a field. An alternative order would be to add copies to the sampled population and then sample that. We follow anatomy: the number of rates received by a field is equal to the number of afferent mossy fibres; the number of *signals* contains duplicate rates.

## 3. The Golgi cell computation

### 3.1 Hypothesis

Despite forming an anatomically seamless carpet of cells, Golgi cells are organised functionally in groups: ensembles. An ensemble is the population of a sagittal row of three fields. This is the minimum unit of the Golgi cell computation. Golgi cell morphology creates ensemble dimensions. The Golgi cell axon ramifies profusely, giving rise to a dense plexus. The axonal field is sagittally elongated – that is, in the direction of the long axis of a microzone (mean range 650 +/- 179 µm by 180 +/- 40 µm [in mice: 33]) – and is the depth of the granular layer, which it fills vertically. Because of the dimensions and orientation of the Golgi cell plexus, all Golgi cells in a sagittal row of three fields inhibit substantially the whole of the middle field. Conversely, each field defines a functional population of Golgi cells – an ensemble – being those cells with convergent input to that field. A single Golgi cell contacts thousands of granule cells [34, 35].

Each mossy fibre terminal is ensheathed by a semi-permeable membrane which restricts neurotransmitter diffusion, a structure termed a glomerulus. Excitation of granule cells and Golgi cells by mossy fibres, and inhibition of granule cells by Golgi cells, all take place here. Fine, beaded axon fibres enter glomeruli, where they inhibit granule cells [34]. ‘In the adult rat cerebellum, each granule cell dendrite receives 2.6 ± 0.55 synaptic contacts from Golgi axon terminals’ [36] citing [32]^1^. However, the large majority of inhibition (98%) is by spillover [4], where neurotransmitter released into the synaptic cleft spills out into the glomerular space. Golgi cells release GABA, an inhibitory neurotransmitter. This is detected by high-affinity GABA_A_ receptors located perisynaptically and extrasynaptically on granule cells [37, 38]. Even synaptically-received signals have most of their effect (i.e., the large majority of charge transfer is mediated) via spillover.^2^

So, an ensemble converts mossy fibre rates received by a three-field row into inhibition of granule cells in the middle field, almost exclusively via spillover. The computation exploits statistical effects called the central limit theorem. Say a population of values (which can have any frequency distribution) is randomly sampled. There are n samples, sample size *m*. If we take the mean (or sum) of each sample, and we take enough samples, the sample means have a normal or near-normal distribution. (This works even if *m* is a range.) The new distribution is centred on the same mean as the sampled distribution, but narrower.

There are three unit populations, each a step in the ensemble computation. First, Golgi cell basal dendrites each quasi-independently randomly sample mossy fibre rates; unit output is a sustained dendritic state which is, at any given instant in time, a linear function of the sample mean. The unit at the second step is the Golgi cell soma; unit output is a linear function of the mean of dendritic states in the physiological form of Golgi cell firing rates, reflecting a sustained somatic state. Third, middle-field glomeruli each quasi-independently randomly sample firing rates of Golgi cells afferent to the field. The effect of the computation is to synchronise inhibition of granule cells in the middle field.

Ensembles conflate sagittally anatomically and functionally. Ensembles in a sagittal strip all return the same output because they all perform the same computation on input data randomly sampled from the same mossy fibre distribution. Unit layers also conflate, so that a unit population extends the length of a rank. The result is to synchronise inhibition of granule cells in the middle field: the Golgi cell population of a rank functions as a single unit.

See section 6.1 for derivation of parameters not given here.

### 3.2 Results

#### 3.2.1 AN ENSEMBLE

Mossy fibre rates sampled by Golgi cells in each field were obtained by sampling from a distribution representing signals received as input to a rank, which were randomly generated subject to constraints on the shape of the distribution, and allowing for mossy fibre terminal branching and clustering, as described in the Methods section (Fig 2 column 1).

**Fig 2.**
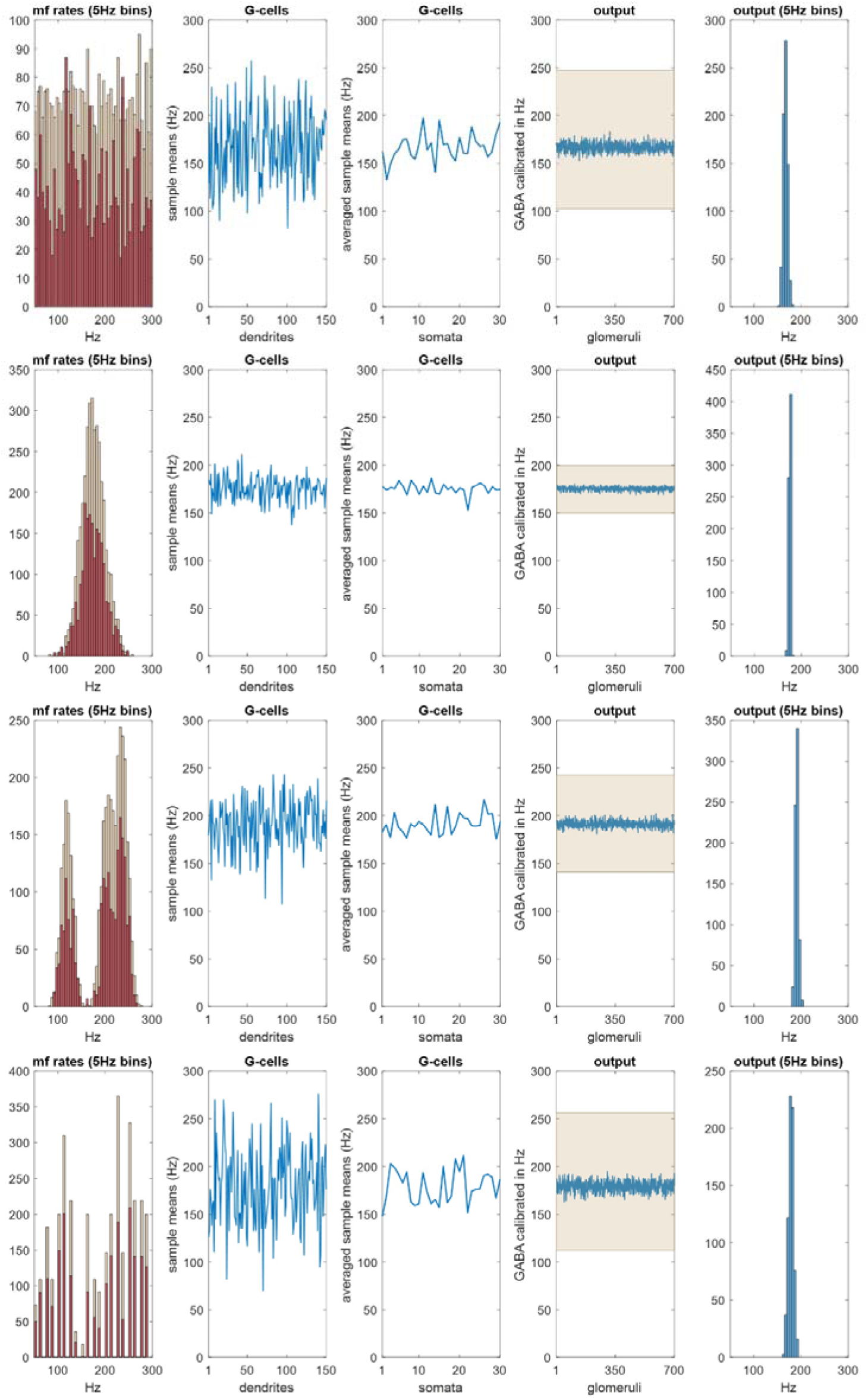
The Golgi cell ensemble conversion. **Column 1** We randomly generated (3,645) values representing mossy fibre rates with a uniform, normal, twin-peaked or discontinuous distribution, simulating input to a row of 100 fields. Rate data were randomly sampled to derive input rates to each field, and copies added to reflect the fact that each cell afferent to a field contributes a randomly variable number of terminals. See Methods for derivation of mossy fibre rates. Pale red: input rates to 100 fields without copies added. Dark red: input rates to a row of three fields with copies added. **Column 2** The first-column dark red data were randomly sampled 150 times, sample size 4, representing mossy fibre input to a 3-field population of 30 Golgi cells, each with 5 basal dendrites (the mean) which each receive contact from 4 mossy fibres. See section 6.1 for derivation of these numbers. Data are the sample means, representing dendritic depolarisation. **Column 3** We next randomly sampled column 2 data *without* replacement 30 times, sample size 5, and took the sample means, representing somatic integration of dendritic signals. **Column 4** Column 3 data were randomly sampled 700 times, sample size 8–12, reflecting convergent input from 3 fields of Golgi cells onto 700 glomeruli in the middle field (which receives the output of a three-field Golgi cell ensemble), and took the sample means, representing intraglomerular GABA concentration, the result of spillover. **Column 5** The column 4 data are presented as a histogram. **All rows** Column 1 mossy fibre data are converted to narrowly-focussed inhibition of granule cells. The mean of the column 4/5 data is linearly related to the mean of the mossy fibre distribution. This result is independent of the shape of the original distribution.

This gave the number of mossy fibre signals, firing rates and the number of copies of each rate received by each field, and therefore input to an ensemble territory, populated by 30 Golgi cells, 10 per field. There are ∼700 mossy fibre terminals per field. As far as we know (and we propose), contact by mossy fibres on Golgi cells is at random at field scale, i.e., signals received by a single basal dendrite are provided by a random sample of local terminals. The number of physiological samplings is equal to the number of basal dendrites per field; sample size is the number of mossy fibres which make contact on each dendrite. The effect is sustained dendritic depolarisation under modulation by a linear function of input rates, which we take as the mean. Sources for this paragraph are cited in section 6.1.

Taking the sample means generates a new set of values (Fig 2 column 2) with a smaller range and centred on the mean, which is linearly related to the mean of the sampled distribution. Basal dendritic charge transfer is passive and little filtered [10, 39]. Integration of dendritic states at the Golgi soma is represented by taking the mean of dendritic charge, for each simulated cell. Golgi cell firing rates have a linear relationship with the amplitude of depolarising current [40], suggesting (and we assume) a linear conversion of somatic membrane potential into the Golgi cell firing rate.

The output is a third data set, Golgi cell firing rates (Fig 2 column 3), on the same normalised scale. Each glomerulus in the middle field receives innervation from a random subset of Golgi cells afferent to the field, and accordingly a random sample of Golgi cell firing rates, mimicking replacement. Golgi cells inhibit granule cells in the glomerulus. A granule cell extends a single dendrite into a glomerulus, typically receiving synaptic contact from two or three Golgi cells [36]. However, inhibition of granule cells by Golgi cells is almost entirely by GABA spillover into the intraglomerular space [4], via extrasynaptic receptors [37, 38], as noted. This effectively increases the Golgi cell to granule convergence ratio.

Golgi cells fire [41] at a time-varying rate in the behaving animal, so that a glomerulus receives continuous input. As a result, there is a sustained build-up of glomerular GABA during behaviour [42] at an adjustable concentration controlled by Golgi cell firing rates [43]. Signalling by spillover is sometimes assumed to be ambient and slow. However, the action of glomerular GABA spillover has a fast phasic component – not as fast as synaptic transmission (∼1 ms) but with a rise time of only a few milliseconds [43]. Unlike the spiky appearance of synaptically-induced inhibitory postsynaptic currents, spillover [3] generates a sustained outward current.

Spillover concentration is under continuous modulation at a level which we take as a linear function of the mean of afferent Golgi cell rates (the sum also works). Sample size is provided by the Golgi cell to glomerulus convergence ratio. The number of Golgi cells afferent to a glomerulus is unreported, as far as we know. We take it as a random number in the range 8–12, generated for each glomerulus. We tested other ratios to confirm that the ratio does not materially affect the ensemble computation (Supplementary Materials).

This generates a final population of values (Fig 2 columns 4 and 5), the output of the ensemble conversion. There are 700 outputs, the number of glomeruli in the middle field. All granule cells that extend a dendrite into the same glomerulus receive equal inhibition (of that dendrite).

#### 3.2.2 ENSEMBLE SUMMARY

There is a progressive narrowing of the data range at each step. Inputs to an ensemble (mossy fibre signals received by a row of three fields) are converted to narrowly-grouped and normally-distributed outputs (representing GABA concentration in middle-field glomeruli). The narrow output range indicates that inhibition is locally synchronised, i.e., granule cells in the middle field receive co-modulated inhibition. Inhibition has a linear relationship with mossy fibre rates, acting through the ensemble conversion. These results are independent of the frequency distribution of mossy fibre rates – input to the system may be in any permutation of firing rates, with the same result.

#### 3.2.3 A RANK

We next used the ensemble conversion to derive inhibition of granule cells in a sagittal row of 100 fields, a rank. The results are shown in Figure 3. As ensembles all perform the same computation and effectively randomly sample (with anatomically simulated replacement) the same data (the distribution of mossy fibre input rates to the whole rank), they all return the same output, or nearly the same output. There are 700 outputs per field, representing glomeruli, as before. The mean of outputs, taken for each field, has a narrow range centred on the mean of mossy fibre input rates to the parent rank. Focus varies with the bandwidth of the sampled distribution (mossy fibre rates). A normal mossy fibre distribution with a range of over 100 Hz is converted to a blue range of ∼3 Hz.

**Fig 3.**
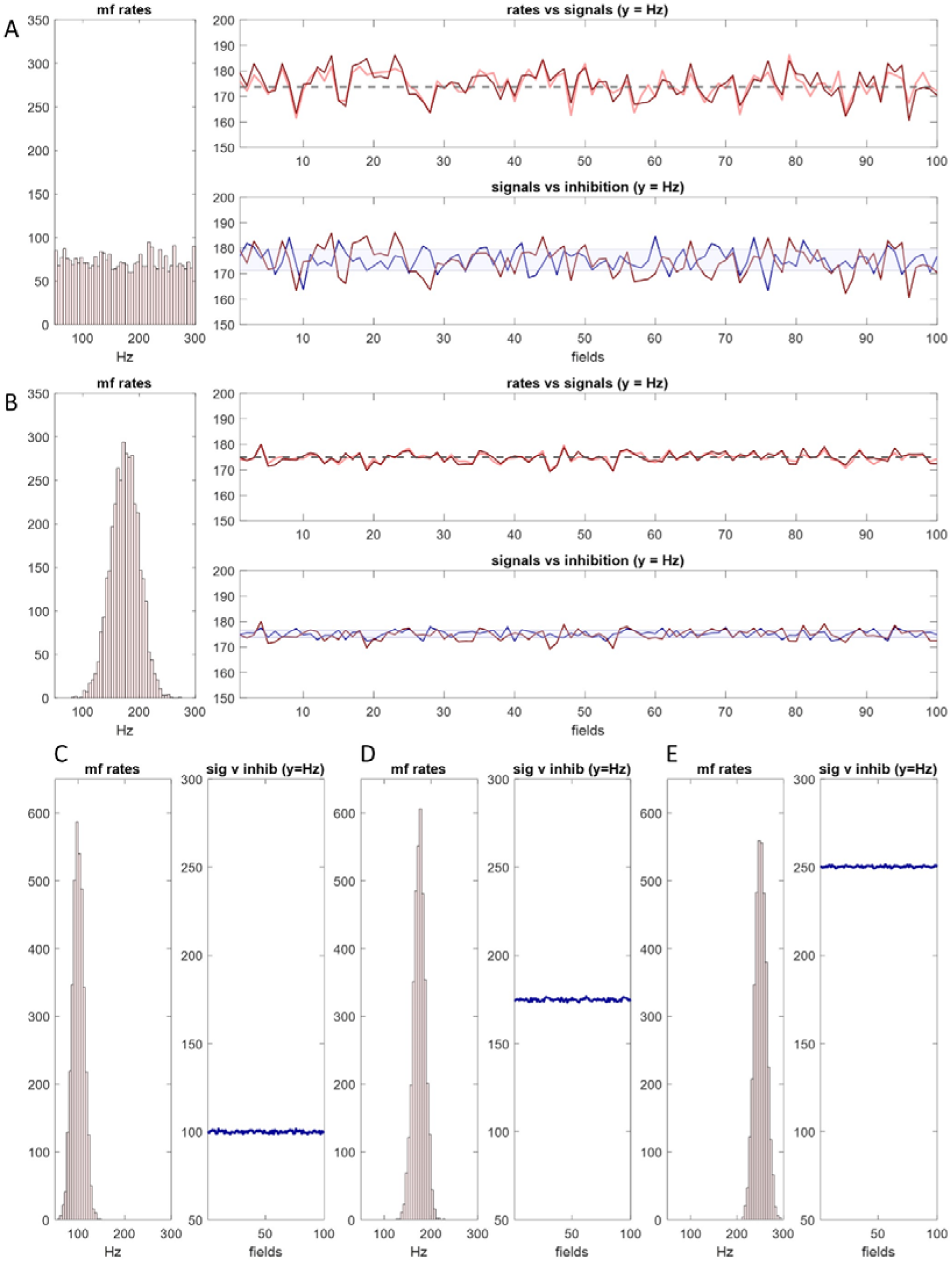
The Golgi cell population of a rank functions as a single unit. **A** Simulation of Golgi cells in a sagittal row of 100 fields, taking the mossy fibre distribution on the left as input (see Fig 2 and Methods for derivation of mossy fibre rates). **Top right:** pink: the mean of mossy fibre input rates to each field; red: the mean with duplicate signals added; dashed grey line: the mean of input rates to the whole rank. **Bottom right:** red: the same as top right; blue: mean, for each field, of glomerulus data derived in the same way as Figure 2 columns 4 and 5 (the ‘inhibitory field mean’); pale blue: SD of glomerulus data. **B** The same as (A) but sampling a different mossy fibre distribution. **A and B** Red data closely track the pink data in top right panels, indicating that duplicate signals (because mossy fibre terminal branches each end in a cluster of terminals) do not skew the mean of firing rates received by a field. Inhibitory field means are tightly grouped: inhibition of granule cells is synchronised in a rank. C The same simulation. Left: the sampled distribution (mossy fibre input rates to a rank); right: the inhibitory field means (we use the means to compare fields). The y axis of the right panel has the same range as the x axis of the left panel. Panels are tall to improve visual comparison. D and E The same, moving the sampled distribution to the right. Inhibition of granule cells is precisely and linearly coupled to the mean of mossy fibre input rates to the whole rank.

In Figure 3 rows A and B, blue data in the bottom right panel represent the mean, for each field, of glomerulus data. The range of the blue data is centred on the grey dashed line – the mean of mossy fibre input rates received by the whole rank – and varies independently of the red data. That is, field-to-field variation of output appears to be inherent in the mechanism, caused by inherent uncertainty of the outcome of random sampling. However, it is not compounded – random variation of the red data doesn’t flow through to amplify random variation of the blue data (in fact the blue data have a slightly smaller range).

#### 3.2.4 RANK SUMMARY

Golgi cells in a rank act as a single functional unit. Ensembles conflate in the sagittal direction functionally as they do anatomically. As a result, the whole granule cell population of a *rank* receive synchronised inhibition which is linearly related to, and varies with, the mean of mossy fibre rates.

## 4. Regulation of granule cells

### 4.1 Hypothesis

Anatomy simulates independent random sampling by a single granule cell of the population of mossy fibre input rates to a rank (section 2). There is a competition for influence between mossy fibre excitation and Golgi cell inhibition of granule cells. The competition takes place in the glomerulus. Granule cells receive input to each dendrite from a single mossy fibre. Granule cells have 3–5 dendrites, averaging 4 (we assume 4, but the result is the same with 3–5). Each dendrite of a single granule cell extends into a different glomerulus. Estimates vary of the number of granule cells that receive contact from a single mossy fibre terminal. 50 is mid-range [31, 32] and consistent with our estimates of the number of terminals and granule cells in a field. Excitation and inhibition compete for influence in a glomerulus. It is unclear if there is always an emphatic winner, but a swing in the dominant direction may be amplified by positive feedback, acting through presynaptic receptors. Activation of presynaptic GABA_B_ receptors on mossy fibres inhibits glutamate release [44], while activation of presynaptic mGluR2 receptors on Golgi cells inhibits GABA release [45]. To fire, a granule cell must receive at least 3 mossy fibre signals which exceed the inhibitory threshold. In experimental conditions, using stimulation that excites very high mossy fibre rates, granule cells can be induced to fire in response to only one input [46].

However, other accounts suggest one is too few [4, 47, 48], and three are needed [49, 50]. We assume 3. Only mossy fibre signals which prevail in the glomerular competition contribute significantly to granule cell depolarisation. During behaviour, the competition is ceaseless. The strength of inhibition is synchronised between glomeruli that populate a rank (section 3), so that all mossy fibre signals compete, at any moment, against the same inhibitory threshold. The outcome in each glomerulus is quasi-independent of the outcome at other terminals, including other terminals that innervate the same granule cell. It is not in fact independent because it is the same at all terminals that arise from the same mossy fibre. However, a single granule cell receives a quasi-independent random sample of outputs of the glomerular competition for the same reason it receives a quasi-independent random sample of mossy fibre rates (section 2). To fire, a granule cell must receive a minimum number of mossy fibre inputs that prevail in the glomerular competition. The probability that enough prevail depends, at any moment, on the strength of inhibition. Since inhibition is synchronised (section 3), the probability is the same, at any time, for all granule cells. The probability that a given granule cell receives three competitive signals and therefore fires is a function of the percentage of mossy fibre signals that are competitive. Expressed as an equation, the expected number of granule cells that fire in a field is the result of

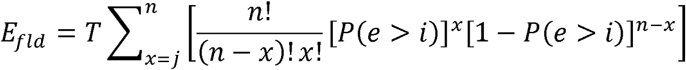

where *T* is the total number of granule cells per field, *n* is the number of dendrites per granule cell, and *j* is the minimum number of dendrites that must receive dominant excitation, *e* > *i*. The proportion of granule cells that fire is near the expected number by the law of large numbers. The percentage of mossy fibre signals that are competitive is always the same because inhibition scales with the mean of mossy fibre rates (because Golgi cells translate mossy fibre firing rates into proportional inhibition). As the percentage is unvarying, so is the probability. This relationship is always conserved regardless of the original distribution (the frequency distribution of mossy fibre rates received as input to the system). So, the probability does not vary either between granule cells or with time. The number of granule cells that fire is very near the expected proportion (the number predicted by probability) because granule cells are numerous. Since the probability is invariant, therefore so is the proportion of active granule cells.

Note: the idea of an all-or-nothing result of the intraglomerular ‘balance’ of excitation and inhibition has appeared previously [51].

### 4.2 Results

The model compares mossy fibre signal frequency with inhibition at each of 4 dendrites of each of 8,750 granule cells per field, in each of 100 fields. Mossy fibre input rates to each field are derived, and duplicate signals added, in the way described earlier. Inhibition – outputs of the ensemble conversion, 700 per field – is provided by the Golgi cell model (which takes the same mossy fibre distribution as input). Mossy fibre signals are randomly sampled. Output of the model is the number of granule cells that fire in each field and the mean of mossy fibre rates they each receive (the sample means). The number that fire in each field is shown in Figure 4.

**Fig 4.**
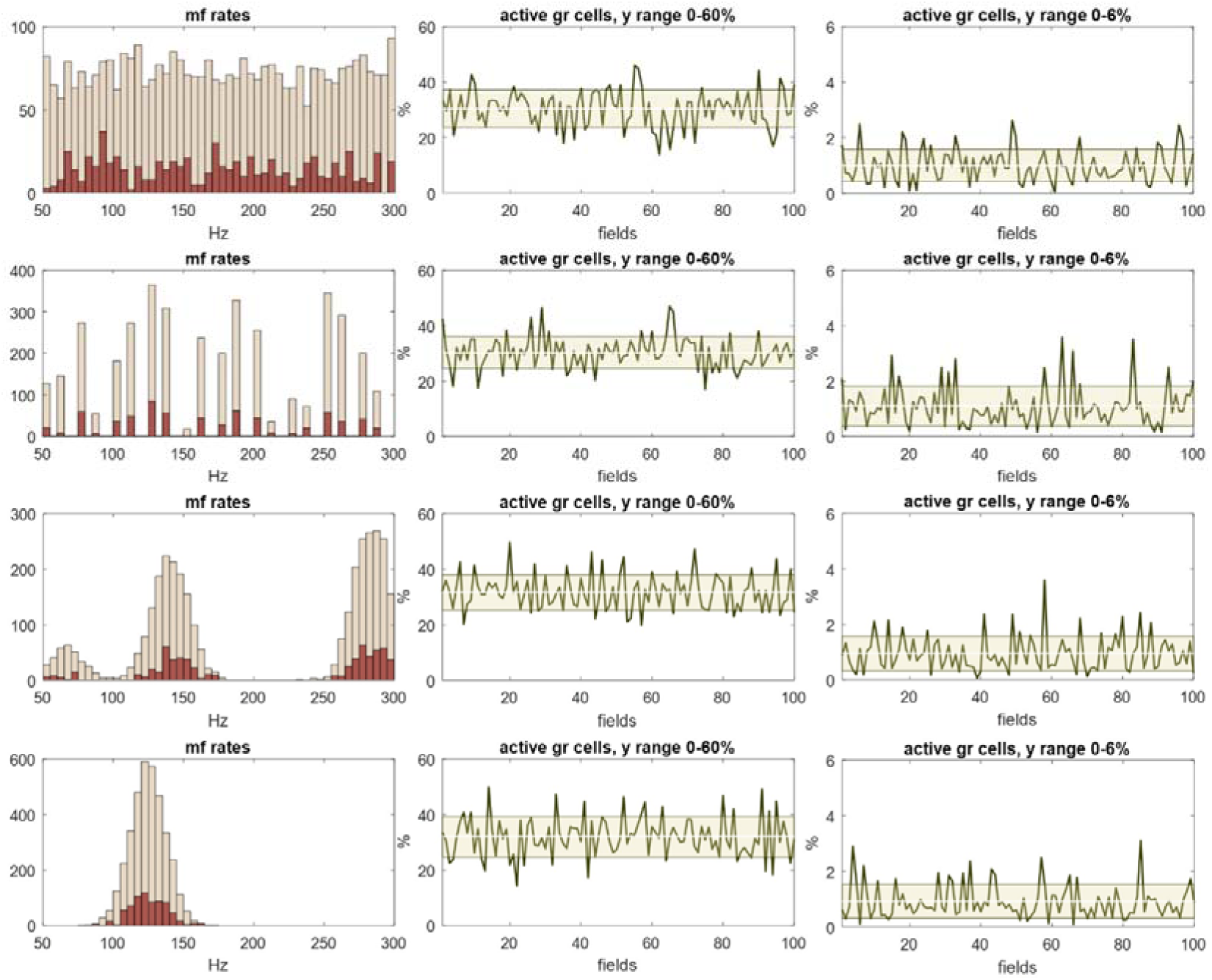
Regulation of granule cell activity by the feed-forward pathway (mossy fibres→Golgi cells→granule cells) We incorporated the Golgi cell ensemble conversion into a simulation of the conversion of mossy fibre signals into granule cell signals (recoding). One of the outputs of recoding is regulation of the percentage of active granule cells. We tested if the recoding computation causes the percentage to be stable, i.e., in the same narrow range in all fields, and whether there is an effect of the distribution of mossy fibre rates. **Column 1** Mossy fibre firing rates were randomly generated with a uniform, discontinuous, three-peak or normal distribution, to represent input to a row of 100 fields. Rates, and rates received by each field, were derived in the same way as earlier Figures. Red data are rates received by a single randomly-selected field, including duplicate signals. y axes are scaled to the data – all distributions contain the same number of signals. **Column 2** The percentage of granule cells that fire in each field. The Golgi cell computation includes no adjustment to sparsen granule cell activity. The white line and shaded area are mean and SD. Regulation is poor: there is wide variation from field to field. **Column 3** The same as column 2 with an adjustment (a co-efficient) to sparsen granule cell activity. y axes have 1/10 the range of column 2. Regulation is strict: the percentage of active cells is confined to a narrow range. This result is independent of the sampled (i.e., mossy fibre) distribution, suggesting that regulation is independent of fluctuation of mossy fibre rates. Nonetheless, field-to-field variation relative to the mean is significant. Homeostatic local fine tuning may be provided physiologically by feedback via granule cell ascending axon contact on Golgi cells, not represented in the simulation.

Feed-forward inhibition – mossy fibre → Golgi cell → granule cell – reliably and closely regulates the fraction of granule cells that are active when the number is small. Variability of the number is inversely proportional to the fraction that are active, suggesting that, if primary regulation is provided by feed-forward inhibition (and we are correct about the mechanism), the physiological number is low – termed sparse code. We would nonetheless expect to see greater precision, suggesting the physiological presence of a supplementary mechanism(s) that we have not represented in the model.

A local feedback mechanism may be provided by granule cell ascending axon contact on Golgi cells. The ascending portion of the granule cell axon makes synaptic contact on local Golgi cells, on basal dendrites [10]. A Golgi cell receives ∼400 functional synaptic contacts from this source, around 25% of the granule cells that a Golgi cell receives contact from in total. During sustained mossy fibre discharge, synapses from local granule cells contribute equivalent charge to Golgi cells as mossy fibre synapses and ‘display similarly fast kinetics’ [52]. Short-latency inhibitory feedback may add homeostatic fine adjustment to feed-forward regulation at local level.

It is a long-standing idea [53] that granule cell activity may be regulated by feedback via parallel fibres, which contact Golgi cell apical dendrites. We have previously proposed that gap junctions between apical dendrites may provide the mechanism for Golgi cells to detect the density of parallel fibre activity [54]. Historically, the feedback pathway – mossy fibre → granule cell → Golgi cell → granule cell – has been regarded as a more intuitive candidate for a regulatory role. It was at one time thought that feedforward inhibition might be involved, instead, in control or adjustment of spike timing. We do not incorporate the gap junction model for these reasons. 1) It has appeared previously. 2) By our calculation, as presented here, the feedforward pathway is the dominant regulator, suggesting the feedback pathway has a supporting, fine-tuning role, so that omitting it has a modest effect on the results. We might expect that feedforward inhibition is the dominant partner because it operates at local (field) level. Strong local regulation is necessary to ensure that active granule cells are uniformly distributed in the general population. The feedback pathway is not sensitive to local fluctuation of granule cell active numbers. 3) Our scope is a single rank. Parallel fibres have a range of around 3mm in each direction from the granule cell soma [29, 30], so that it would be necessary to model a larger region (about 40 times the size), which extends into the molecular layer, to generate parallel fibre input to a strip of Golgi cells. 4) It is a different (and independent, although related) mechanism. 5) They have (related but) different functions. The feed-forward pathway is a local regulator of the fraction of active granule cells (with an effect that flows through to parallel fibres), whereas the feedback pathway is a regulator of the fraction of active parallel fibres only. While the feedback mechanism works by modulating inhibition of granule cells, modulation is very weakly related to local conditions.

Uniform density of active granule cells, and therefore strong local regulation, is important for mossy fibre information to be faithfully coded by granule cells. The granule cell code is the subject of the next section.

The results are independent of (i) the frequency distribution of mossy fibre rates; (ii) whether mossy fibres are all active or some are active and some are not; and (iii) the permutation they are active in (which ones are active and which are not).

## 5. The granule cell code

### 5.1 Hypothesis

Granule cells code information in their collective activity, in functionally defined groups. Granule cell code groups are functionally defined by the topography of mossy fibre input to the cerebellum, as Golgi cell networks are. The output of a rank is the smallest unit of granule cell encoded information. Granule cell encoded information, accordingly, has spatial dimensions, the dimensions of a code group. Information is coded in the frequency distribution of granule cell firing rates. Therefore, the same information is coded, at any moment, in firing of any random sample of co-active cells, provided the sample is large enough. Because granule cells are very numerous, the minimum dimensions of code are much smaller than a code group, because they are the size of dimensions that are large enough to contain a sufficient number of co-active cells at the regulated level of granule cell activity. The permutation of active mossy fibres, and of granule cells, and how firing rates are distributed among active cells, all code nothing.

Anatomy simulates independent random sampling by a single granule cell of the population of mossy fibre input rates to a rank (section 2). The sampling unit is the whole cell since it receives only a single input rate to each dendrite. Sample size is therefore 4 (on average), and the number of samples per field is the number of granule cells, which we estimate at 8,750 (see Methods). The unit function is only operative in the case of a cell that receives at least the minimum number of competitive mossy fibre signals. Then, somatic depolarisation is a linear function of the mean (or sum) of competitive input rates (and is in turn straightforwardly related to the granule cell firing rate – see section 6 for a discussion and references). We postulate that the linear input-output relationship is undiluted by inhibition because dominant signals displace a significant effect of inhibition. Derived in this way, there is a reliable statistical relationship of granule cell rates and mossy fibre rates, again by the central limit theorem. The range of granule cell rates is narrower than the range of mossy fibre rates, centred on the mean, and attracted to a normal distribution. The mean of the new distribution (unit outputs) is linearly related to the mean of mossy fibre rates. This result is repeated in all fields. As all fields randomly sample, at any time, the same mossy fibre distribution, and perform the same computation, they all return the same granule cell code. This provides the substrate for granule cells to collectively encode mossy fibre rate information.

### 5.2 Results

We ran the same simulation as section 4 to test whether, and how, mossy fibre rates are related to granule cell rates, and if fields in a rank all have the same output. As before, the model compares mossy fibre rates with inhibition at each of 4 dendrites of each of 8,750 granule cells per field, in each of 100 fields. Mossy fibre signals received as input to each field are derived in the same way, described in section 2, and inhibition is provided by the Golgi cell model. The results are shown in Figures 5 and 6. We stated in section 4 that the outputs of the model are the number of granule cells that fire in each field and the mean of input rates they each receive – the sample means.

**Fig 5.**
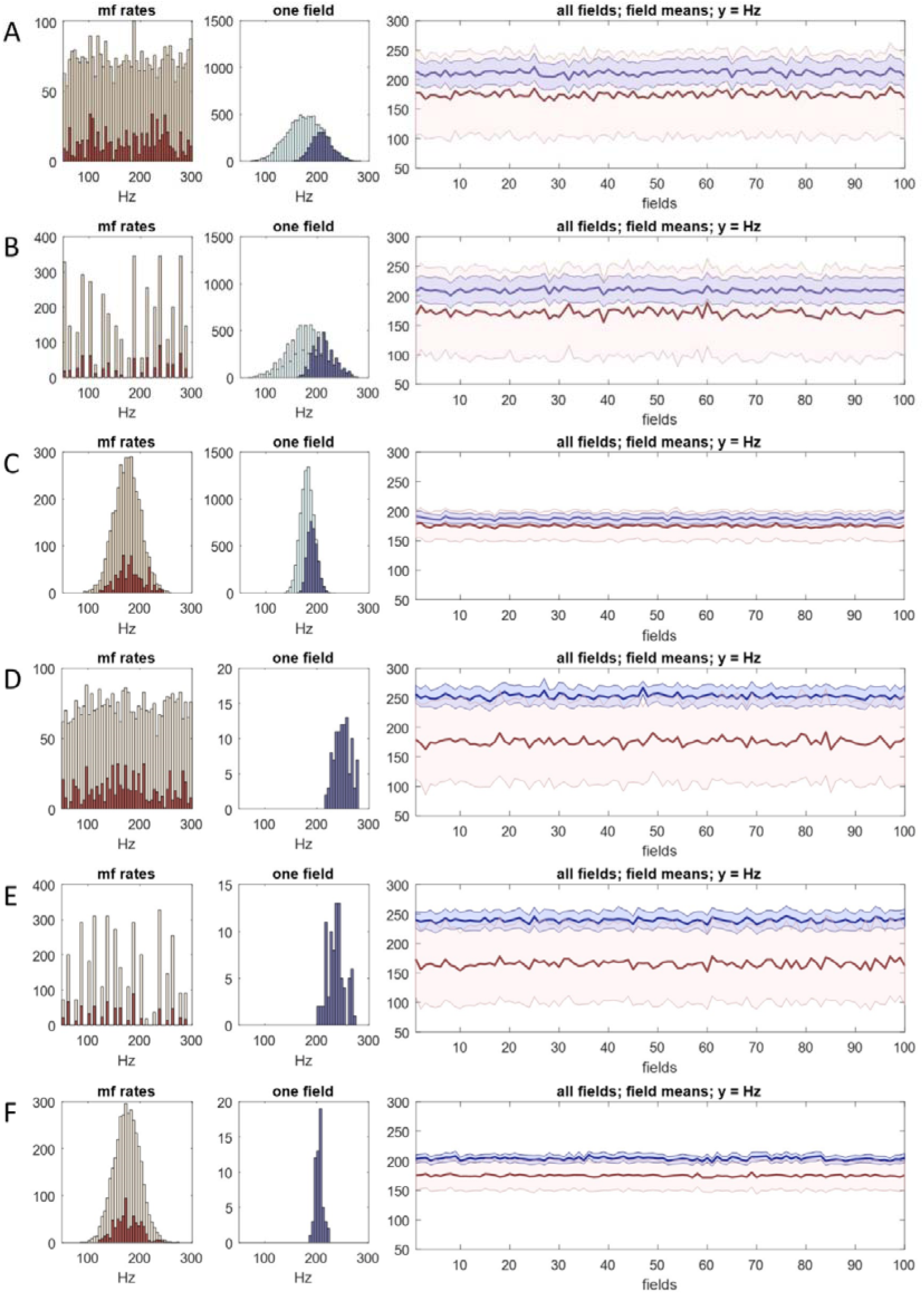
A rank functions as a single unit. The same simulation as Figure 4 but data are granule cell firing rates or derived from granule cell firing rates. **Column 1** Mossy fibre rates were generated in the same way as earlier figures (the number of signals is the same in all rows). **Column 2** Pale green: sample means – the mean of input rates to each granule cell in a single field selected at random; blue: sample means of the subset of granule cells which meet the conditions to fire. In rows A–C, ∼30% fire. In rows D–F, ∼1% fire, y axes are rescaled and green data are omitted. **Column 3** Red: mean of mossy fibre input rates to each field in a rank, including duplicate signals; pink: SD of mossy fibre input rates to each field; blue: field means (i.e., the mean of granule cell firing rates in each field); pale blue: SD of granule cell firing rates in each field. Regardless of the shape and large range of the mossy fibre distribution, the field means always converge towards a straight line which is a fixed distance from the overall mean of mossy fibre rates. The amount and direction of field-to-field random fluctuation of the blue data is independent of local fluctuation of the red data, and in a smaller range. Uncertainty inherent in random sampling – first by single fields of input rates to a rank, and then by Golgi cells and granule cells of input rates to a field – is not compounded.

**Fig 6.**
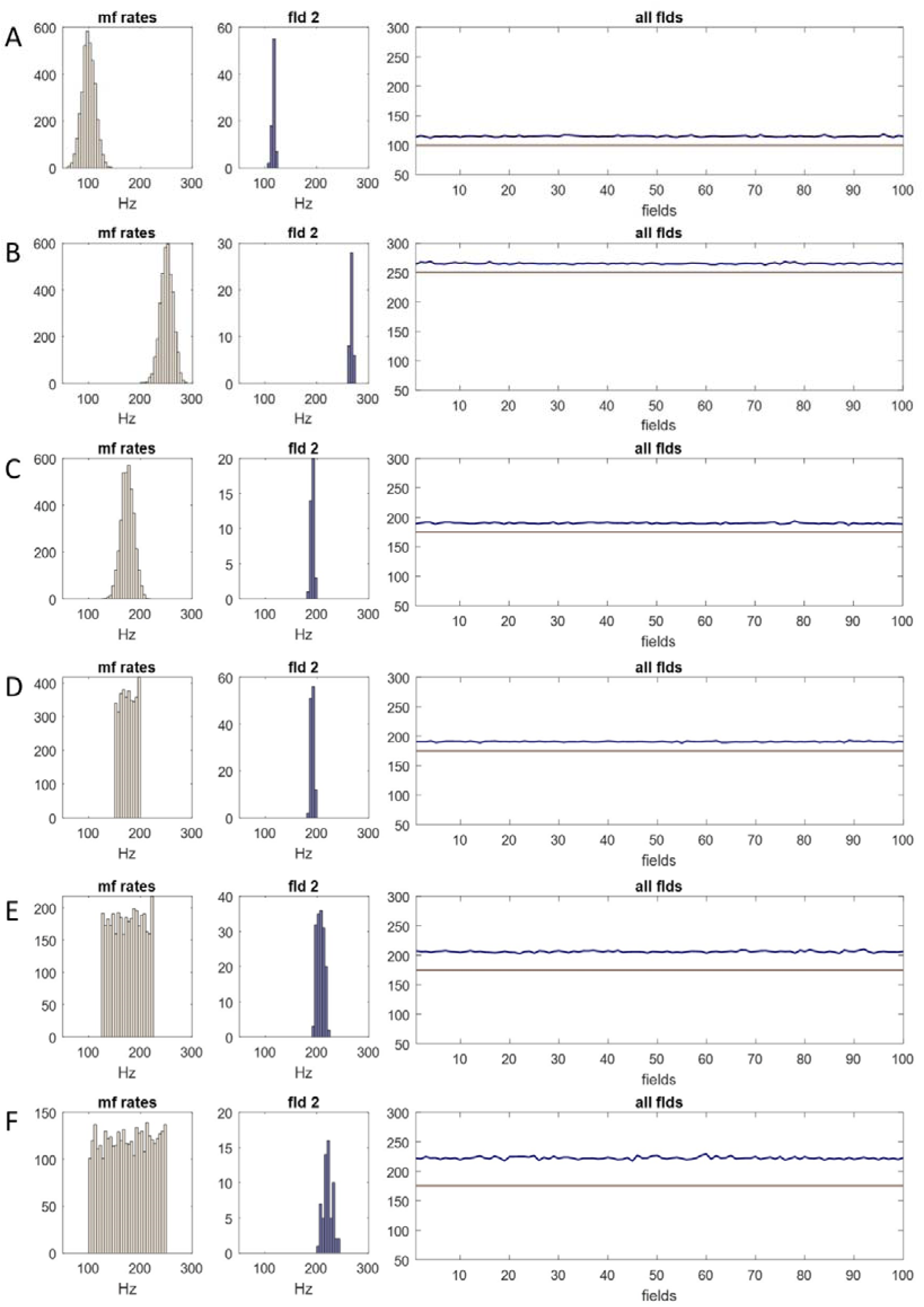
Granule cells collectively code mossy fibre rate information. **Column 1** Mossy fibre rates were generated in the same way as earlier Figures, with either a normal (SD 12.5) or uniform distribution, representing input to a row of 100 fields, with copies added – the sampled distribution. **Column 2** Histogram of granule cell firing rates in a randomly selected field, obtained in the same way as in the bottom 3 rows of Fig 5. **Column 3** Grey line: the mean of the sampled distribution; blue line: field means. **A–C** The fields means are precisely and linearly related to the mean of mossy fibre rates. Blue data approach a straight line which is parallel to the grey line at a mean distance of 15 Hz (SD 1.21 Hz). **D–F** If the shape and mean of a uniform sampled distribution are unchanging, the distance between the blue and grey lines is linearly related to the range of the sampled distribution.

The narrowing and predicted shape of the granule cell frequency distribution is evident in all conditions. The ‘field means’ (for each field, the mean of the sample means) approach a straight line, the test that a rank operates as a single functional unit. When mossy fibre rates are normally distributed (SD 12.5 Hz), the granule cell distribution is very narrow (and normal if rates are counted in narrower bins) and the field means are very close to a straight line. The field means follow the mean of mossy fibre rates at a fixed distance. In all tested conditions, granule cell rates have a narrowed range, which is the same in all fields, clustered around the mean, and attracted to a normal distribution (more evident with narrowly and/or normally distributed mossy fibre rates). This holds even though a single granule cell receives a tiny fraction of mossy fibre rates, and each field receives only a small fraction of mossy fibre signals received by a rank, and also regardless of the shape of the mossy fibre distribution.

What may we infer from that? The purpose of the demonstration is to show that a network of linear units, in the form described, has the predicted output. We know (or can derive) from reported anatomy the size of unit populations, population ratios, and convergence and divergence ratios. Evidence is cited in context. The evidence for linear units is discussed in section 6. Random sampling is a functional feature, a proposal to account for anatomy.

Predictions confirmed: input rates to the system in the physiological range, with any distribution and incorporating duplication as a result of terminal branching and clustering, in anatomical detail including variation of single cells, is reliably converted to a granule cell distribution with the predicted features. A key feature is that inhibition of granule cells is synchronised both in a field and in a rank. A second key feature is that mossy fibre rates are converted to a granule cell distribution with the same shape and mean in all sectors of a rank, so that any random sample of granule cell rates also has that distribution.

The principal outstanding questions, as we see it, are whether:

1. Computational units are linear (see next section).
2. Contact between cells in the granular layer is at random below a scale threshold set by range and morphology.
3. Sampling mimics replacement, an inference from anatomy.
4. The frequency distribution of granular layer firing rates is code.

If so, we can go further. Granule cell firing conserves mossy fibre rate information, coded at the collective scale of the output of a rank. This is therefore the minimum resolution of granule cell information. However, it is not the minimum unit of code. The same information is coded in any random sample of granule cell signals, over a minimum number. It is immaterial (i.e. codes nothing) which particular cells are active or which cells fire at what rates, or even what rates those are.

## 6. Support for linear transmission

There is a growing body of evidence of linear communication in the granular layer, indicating that unit functions are linear. This section, except the first subsection, is a short review.

### 6.1 Derivation of some Golgi cell ensemble parameters

4 Purkinje cells fit in a field, based on field size, thickness of the Purkinje cell dendritic plate – 9 µm [55] – and the distance between them – ∼40 µm [56]. ‘In man, the ratio [of Purkinje cells to Golgi cells] is 1∶1.5, while in the monkey and cat it is almost 1∶1.9 and in the rat 1∶3.3’ [57]. Possibly, estimates are low because smaller Golgi cells at deeper level were not counted [56 p.121]. We take it that there are 10 Golgi cells per field, a ratio of 1:2.5 with 4 Purkinje cells.

Contact by mossy fibres on a Golgi cell is on basal dendrites. 4–6 basal dendrites arise from the Golgi cell soma. We assume 5, so a population of ensemble-grouped Golgi cells has a combined total of 30 x 5 = 150 basal dendrites. The number of mossy fibres which contact a Golgi cell is unreported. Dendrites radiate from the cell body. A single dendrite has a range of 50–100 µm [56]. The longest dimension of a field – which contains ∼700 mossy fibre terminals – is 200 µm. We estimate the mean number of mossy fibres that contact a single dendrite is 4, so 20 per cell, or ∼3% of the population of terminals (or 0.6% per dendrite).

Glomeruli individually receive innervation from a random sample of Golgi cells afferent to a field. It is unknown how many Golgi cells innervate a glomerulus.

### 6.2 Linear transmission of mossy fibres to Golgi cells

#### 6.2.1 MAIN DISCUSSION

Golgi cells are large interneurons whose cell bodies and basal dendrites lie in the granular layer. Mossy fibres directly contact Golgi cell basal dendrites. Golgi cell firing frequency increases linearly with the amplitude of depolarising current [40 p.845]. ‘Golgi cells can follow peripheral signals in a continuous fashion, modulating their frequency with the intensity of the stimulus [citing [58, 59]]’ [60]. As a result, during movement, the ‘output of Golgi cells varies smoothly as a function of input.…They are capable of modulating their output to reflect their input with little noise or signal distortion’ [11]. ‘Sensory-evoked Golgi-cell inhibition scales proportionally with the level of mossy fiber excitatory synaptic input’, such that inhibition reliably conveys mossy fibre rate information [3]. On this evidence: mossy fibre rate information is conserved proportionally in Golgi cell firing rates.

Golgi cells extend their basal dendrites into glomeruli [34, 61]. Contact on them by mossy fibres is multi-synaptic [56], contributing to a reliable, rapid (submillisecond) response. The fast rise time and weak distance dependence of the amplitude and timing of Golgi cell EPSCs evoked by mossy fibre signals, suggests that basal dendrites introduce relatively little filtering [10, 39], as expected for large-diameter dendrites. We note that the pause in Golgi cell firing following a discrete stimulus under anaesthesia [3, 62, 63] disappears from Golgi cell recordings during locomotion [59].

During behaviour in freely-moving animals, mossy fibre activity is dense: a high fraction of mossy fibres are active [64]. Both mossy fibres [65–67] and Golgi cells [59] have been reported to fire with a sustained, time-varying signature. In the behaving animal, Golgi cell basal dendritic membrane potential is a continuous variable under modulation by sustained inputs, we submit. As charge transfer to the soma is passive, polarisation of the soma is likewise sustained, under modulation by dendritic states.

It is worth noting that very high mossy fibre rates can be generated in experimental conditions. However, the typical physiological range is 50–300 Hz [66]. In the simulations, we use the physiological rates, within stated constraints on the shape of the distribution.

#### 6.2.2 IS THERE OTHER CONTROL OF GOLGI CELLS?

Do Golgi cells receive a significant influence of input from other sources? The evidence is incomplete but currently suggests other influence is absent or weak. Contrary to early reports, neither Purkinje cells nor climbing fibres contact Golgi cells [68, who give references], and only a modest minority of inhibitory inputs to Golgi cells (if any) are from molecular layer interneurons [69], which generate weak synaptic currents [70], consistent with either extremely weak or wholly absent innervation [71]. There is conflicting evidence whether Golgi cells make inhibitory synaptic contact on each other [71, 72]. Our simulation does not include an effect on Golgi cell firing by other sources of input.

#### 6.2.3 GOLGI CELL OSCILLATIONS

Golgi cells fire autonomously. Under anaesthesia, firing falls into a slow oscillating pattern [73]. Under excitatory synaptic input, however, this disappears [72]. Anaesthesia ‘has a strong influence on spontaneous activity of Golgi cells’ [68]. Discharge at low rates with irregular timing under anaesthesia [41] is replaced by higher rates with more regular timing without anaesthesia [58, 59, 74]. This paragraph is a cautionary note about evidence obtained under anaesthesia, which has been used to argue that oscillations provide a form of signalling.

### 6.3 Linear transmission of mossy fibres to granule cells

There is significant evidence that granule cell firing rates are linearly related to input rates, but also evidence that has been given a conflicting interpretation. We take them in turn.

The short and equal length of granule cell dendrites would suggest light and equal filtering. The mossy fibre-granule cell connection has a range of adaptations which support high-fidelity transmission of high frequency signals across a wide bandwidth [8]. Faithful transmission of rate information is reported [1, 46]. Vesicle release and replenishment are fast [9, 12] and postsynaptic AMPA receptors operate in their linear range [12], where they are resistant to desensitisation [75]. Multiple contacts are made by a mossy fibre with each granule cell [32]. Conversion of depolarising somatic charge into granule cell firing is a simple function, such that granule cells ‘have a relatively linear and uncomplicated conversion of depolarisation level to spike rate’ [48 p.2393, citing Jorntell and Ekerot 2006 and D’Angelo et al 1998].

Glutamate spillover enhances precision and reliability of transmission [12, 76]. During behaviour, at sustained physiological mossy fibre rates, there may be a build-up of intraglomerular glutamate, increasing the relative influence of spillover in a balance with synaptic transmission, assisted by short-term synaptic depression. In these conditions, spillover may dominate, as it dominates inhibitory transmission [4, 43]. Spillover from multiple release sites (200–400 at a single mossy fibre terminal) is plausibly sufficient to mitigate variability of vesicular size, release probability, and the number of release points at synapses made on any individual cell, increasing fidelity and equality of excitation of granule cells.

Against that, there are reported to be heterogeneous mossy fibre-granule cell synaptic weights. Different strength and short-term dynamics of the mossy fibre to granule cell connection have been reported in vestibular slices, with GABA_A_ receptors blocked. The amplitude of the response, measured at the soma, is reported to be statistically correlated to the source of the mossy fibre that is stimulated [77], suggesting that the strength of the connection depends on the source.

However, the presence of weights does not necessarily mean that the function is to modulate firing rates. Rather, the authors themselves suggest that the function may temporally code the source of inputs because different combinations have different response onset times. An alternative, which they also mention, is that some inputs may be in a supporting role, reminiscent of ‘driver’ and ‘modulatory’ signals in the thalamus and cortex [78]. Feasibly, signals in a supporting role must be present for the postsynaptic granule cell to fire, but do not affect the granule cell firing rate.

Heterogenous granule cell subtypes have been inferred from recordings of granule cells in rat slices, from the finding that when stimulation is extended beyond 500 ms firing of most cells changes, either slowing down or (in fewer cases) speeding up.

However, the significance is not clear. Granule cells typically fire in short bursts – shorter than the duration of the stimulation tested, so perhaps not long enough for the effects observed. Short granule cell burst duration may be because a granule cell normally does not meet the conditions to fire for long, because few fire at any time, and the permutation of active and inactive inputs to a single granule cell, and the firing rates of active inputs, are both in a state of restless change. Also, excitation of granule cells receives a concurrent regulatory influence of a network of Golgi cells which populate a region which receives 100s of time-varying mossy fibre signals at any time (so, not the conditions tested). Confirmation of functional significance of adaptive firing in experimental conditions is outstanding.

### 6.4 Linear transmission of Golgi cells to granule cells

The hypothesis includes the proposal that inhibition of granule cells is, at any time, linearly related to the mean firing rate of Golgi cells afferent to each glomerulus. In support, we cite evidence that GABA spillover into the intraglomerular space is proportional to afferent Golgi cell rates and that inhibition of granule cells is almost exclusively mediated by spillover, and therefore linearly reflects the intraglomerular concentration of GABA. Citations appear in the Golgi cell section but for convenience they are repeated here.

Fine, beaded axon fibres enter glomeruli, where they inhibit granule cells [34]. ‘In the adult rat cerebellum, each granule cell dendrite receives 2.6 ± 0.55 synaptic contacts from Golgi axon terminals’ [36] citing [32].^3^ However, the large majority of inhibition (98%) is by spillover [4], where neurotransmitter released into the synaptic cleft spills out into the glomerular space. Golgi cells release GABA, an inhibitory neurotransmitter. This is detected by high-affinity GABA_A_ receptors located perisynaptically and extrasynaptically on granule cells [37, 38]. Even synaptically-received signals have most of their effect (i.e., the large majority of charge transfer is mediated) via spillover.^4^

Golgi cells fire [41] at a time-varying rate in the behaving animal, so that a glomerulus receives continuous input. As a result, there is a sustained build-up of glomerular GABA during behaviour [42] at an adjustable concentration controlled by Golgi cell firing rates [43]. Signalling by spillover is sometimes assumed to be ambient and slow. However, the action of glomerular GABA spillover has a fast phasic component – not as fast as synaptic transmission (∼1 ms) but with a rise time of only a few milliseconds [43]. Unlike the spiky appearance of synaptically-induced inhibitory postsynaptic currents, spillover [3] generates a sustained outward current.

## 7. Discussion

### 7.1 Summary

Network models are often a simplified representation of physiology to provide a substrate for sophisticated computations. We propose the reverse: physiology is highly adapted to provide a substrate for straightforward computations. In summary we propose:

1. There is a computational effect, unaided and unlearned, of the combination of cell morphologies, network architecture and linear units, for which the granular layer is highly adapted.
2. Physiology is the physical form of a network of linear units organised in layers. Units randomly sample afferent activity. Code groups communicate by a facsimile of independent random sampling. Anatomy mimics random sampling with replacement by a single cell of the entire population of firing rates in the afferent cell layer.
3. Data processing exploits statistical effects of random sampling, such that i) inhibition of granule cells in a rank is synchronised and tracks mossy fibre rates; and ii) the frequency distribution of granule cell rates is reliably (because statistically) well-predicted by mossy fibre rates.
4. High-dimensional mossy fibre information recodes as a very low-resolution granule cell code. Large granule cell groups function as a single unit, coding information in their collective activity.
5. At any instant, a whole group codes the same information. Accordingly, granule cell-coded information has spatial dimensions, the dimensions of code groups. As code groups are long and thin, information has a striped pattern viewed from the cerebellar surface.
6. Code groups span the cerebellar network in the sagittal direction, from side to side, mirroring microzonal organisation of the molecular layer. Like microzones, they are defined topographically, by their input from outside the cerebellum, rather than having physical boundaries (but they have fuzzier borders).
7. While firing of a whole code group is the minimum unit of information, it is not the minimum unit of code. The same information is coded in any random sample of code-grouped granule cell firing rates. In that sense, the code is homogenised below a scale threshold.
8. The shape of data and homogenised code are key to function, having results discussed below.

### 7.2 Commentary

#### 7.2.1 GENERAL COMMENTARY

Speaking generally, there is a conflict between evidence relating to learning and evidence of linear transmission. There is a conflict partly because there is an implicit inference that i) synaptic changes are functional, and ii) synaptic weights are graded. Historically, there has been strong emphasis on researching learning [79, 80]. The traditional learning model, which centrally features synaptic memory, has not received confirmation by experimental work. However, there is still a mainstream feeling that the computation will turn out to involve synaptic weights. In this view, core assumptions are retained, and the aim of further work is to understand how plasticity at multiple sites is related, and related to function.

Such has been the dominance of learning, it is regarded as naive to have a different opinion. However, it does not address a range of evidence relating to linear transmission (discussed in context, in the main text, and reviewed in section 6), and of linear relationships between firing rates and behavioural metrics, suggesting linearity is not merely an experimental artefact (for which Raymond and Medina provide 30 references [15]). We present an alternative approach where the focus is to explain cell morphologies/general anatomy and linear transmission. We propose there is a computational effect of anatomy, give linear signalling. Our focus is the granular layer of the cerebellar cortex. We deal with the linear/non-linear question by dedicating a section (section 6, as noted) to a review of the evidence for linear transmission, which is rich for the granular layer.

We know of course that this is contentious. However, it has merits.

1. It has in fact good explanatory value – cell morphologies, granular layer network geometry, cell ratios, spatial organisation, the shape and size of dendritic and axonal territories, convergence and divergence ratios, random contact that is on some and not other cells that lie in axonal range, the duplication of mossy fibre signals as a result of terminal branching and also locally because branches each end in a cluster of terminals, the structure and physiology of the glomerulus, and linear signalling (including evidence that synaptic connections are adapted for high-fidelity transmission) – are all accounted for.
2. We propose – as artificial neural network models do – that signals traffic flows through computational units organised in layers, but with the difference that

i. Our units are matched with physiological hardware;
ii. Units form functional populations whose size and spatial dimensions are an effect of known physiology; and
iii. Unit functions are not conjecture, they are linear, with the basis in evidence of linear signalling. We note in this connection that non-linear synaptic properties predicted by experimentally observed synaptic changes have the same problem with evidence of linearity, but reversed, as linear unit functions do with synaptic evidence of non-linearity.
3. A linear network, as we describe, has functional advantages. We discuss the functional significance of the proposed form of the code, and of the computation, in the remainder of the Discussion. We make three main arguments. In all three, the form of the computation solves a problem. The first problem is how an anatomically seamless tangled mat of cells is organised functionally with a view to convert high-dimensional external signals into coherent firing of microzone-grouped Purkinje cells. The second is that – against expectation – the pattern in which parallel fibres are active may not in fact be reliably or perhaps ever replicated. Theory has previously addressed this problem by making assumptions that have been widely adopted but which are not safe. The first paragraph of the paper comments on this. The third problem is how insentient cerebellar circuits are able to process input data usefully with, as far as we know, little or no information about where it is from or what it represents.

#### 7.2.2 CONTROL OF MICROZONE-GROUPED PURKINJE CELLS

There is a huge sacrifice of data resolution on entry to the cerebellum, counter to the general expectation that data processing is sophisticated. This is functional. Coded in this form, granule cell data can drive coherent firing of large cell groups downstream. Microzones lie at right angles to parallel fibres and parallel to code groups. The statistical form of the granule cell code means the same information is received by all locations of sagittal plane, so that microzone-grouped Purkinje cells can receive the same information without receiving any of the same signals. An estimated 350,000 parallel fibres pass through a Purkinje cell dendritic territory [28, 81–83]. If even only a small fraction of parallel fibres is active (say, 0.5%), and allowing both for the fact that only one in two make contact and that a majority of synapses are severely depressed or silent [84, 85], a single Purkinje cell still receives around 200 parallel fibre signals at working synapses at any time. A granule cell code in this form provides a means to modulate firing of microzone-grouped Purkinje cells in unison, ultimately under control of mossy fibre rates. There is evidence that Purkinje cells linearly code the strength of granule cell inputs [5–7] and linearly code behavioural metrics [2, 86–89].

#### 7.2.3 REPLICABILITY

A statistical code may be a physiological strategy to mitigate variable performance of single neurons and synapses, long known to be a practical problem for replicable biological information capture. The traditional cerebellar learning model [53, 90, 91] relies on (and assumes) replicability because pattern replication is necessary for signals to match up with the correct training-modified parallel fibre synaptic weights. Learned changes of synaptic weights are also a core feature of many neural network learning models, not just cerebellar models.

However, this may not be safe. First, it is not clear that input to the cerebellum is ever replicated. Around 4,000 mossy fibres innervate the region that supplies parallel fibre input to a single Purkinje cell. It may not be a safe assumption that conditions upstream are routinely repeated with precision sufficient to consistently (or perhaps ever) generate a repeat of the same inputs. Second, single neurons and synapses are not reliably consistent performers. Neurons even of the same type are by no means replicas, and synaptic transmission involves random variables. Third, the granular layer is *designed* so that small differences of input are converted by decorrelation to wholesale changes to output (the pattern of active parallel fibres). Indeed, even that is not necessary – the same pattern of active mossy fibres but firing in a different permutation of rates also generates a different pattern of active granule cells, and therefore parallel fibres.

Homogenised granule cell/parallel fibre code removes the problem because replication of the pattern of active granule cells is unnecessary to precisely code mossy fibre information. This is because:

1. It is coded in granule cell firing rates and not the pattern they are active in.
2. It is coded in population activity, in rate statistics.

In the traditional model, a granule cell rate code was effectively redundant. That is not to say granule cell rates had no effect on Purkinje cell firing. But the function of synaptic memory was to *displace* control by rates. They might start out as *any* rates, because learning would generate the ‘desired’ firing of Purkinje cells following training. In the alternative model, output of the system is an unlearned response to input rates and the pattern of active cells is irrelevant. The colossal sacrifice of resolution is a functional feature.

#### 7.2.4 WORKING ‘BLIND’

A perennial issue for network theory is that biological networks do not seem to have much information about the signals they receive. Input to the cerebellum is eclectic and in multiple modalities. In some cases, a single granule cell can receive multi-modal input. In the traditional cerebellar model, supervision solves this problem. It is unnecessary for the cerebellum itself to discriminate because it is an assumption of the model that externally-sourced instruction signals train the correct response.

We propose instead that the solution is to sacrifice the resolution of high-dimensional input data. Source and modality data are disregarded below a topographically-defined scale threshold. It is therefore unnecessary to discriminate below that scale. Mapping studies show that mossy fibre signals (evoked by cutaneous stimulation) share the topographical organisation of climbing fibres [19, 92, 93]. At sub-topographical dimensions, we submit, signal source and type are simply discounted. Therefore, networks do not need to be adapted to the source or type of input they receive – anatomically, networks are interchangeable, so network design can be modular. This feature allowed basic cerebellar wiring to be highly conserved through dramatic upheavals of the vertebrate phenotype [94, 95].

## Glossary

**field** A nominal division of the granular layer measuring 200 µm (sagittal) x 150 µm (mediolateral), the average dimensions of the area enclosing a mossy fibre terminal cluster.

**field mean** For each field, the average of the mean of mossy fibre input rates to each granule cell which meets the conditions to fire.

**rank** A sagittal row of 100 fields.

## Grants

Funding: This work was supported by the Leverhulme Trust [grant number ECF-2022-079] (Mike Gilbert); the Swedish Research Council [grant number 2020-01468]; the Per-Eric och Ulla Schybergs Foundation [grant number 42630]; and the Crafoord Foundation [grant numbers 20180704, 20200729, 20220776, & 20230655] (Anders Rasmussen).

## Acknowledgement

Dr Mike Evans, School of Mathematics, University of Leeds, LS2 9JT, UK, read an early draft of the paper and provided feedback.

## Author contributions

M.G. Researched, conceived and developed the ideas, performed the analysis, wrote the code, prepared the figures and wrote the manuscript. A.R. provided expertise and feedback, and reviewed and provided editorial comments on the manuscript.

## Declaration of interests

The authors declare no competing interests.

## Footnotes

1 Data are sparse. The original data are an estimate derived from visual inspection of slices thirty years ago so far without later corroboration, except from Rossi and Hamann 1998, who derive a lower estimate but propose the disparity is because their estimate is too low.

2 Because of the long duration of IPSCs generated by spillover (recorded in slices), the total charge carried is three times that of IPSCs generated by directly connected terminals. 36. Rossi, D.J. and M. Hamann, *Spillover-mediated transmission at inhibitory synapses promoted by high affinity alpha6 subunit GABA(A) receptors and glomerular geometry*. Neuron, 1998. **20**(4): p. 783-95.

3 Data are sparse. The original data are an estimate derived from visual inspection of slices thirty years ago so far without later corroboration, except from Rossi and Hamann 1998, who derive a lower estimate but propose the disparity is because their estimate is too low.

4 Because of the long duration of IPSCs generated by spillover (recorded in slices), the total charge carried is three times that of IPSCs generated by directly connected terminals. 36. Rossi, D.J. and M. Hamann, *Spillover-mediated transmission at inhibitory synapses promoted by high affinity alpha6 subunit GABA(A) receptors and glomerular geometry*. Neuron, 1998. **20**(4): p. 783-95.

## Notes

### Competing Interest Statement

The authors have declared no competing interest.

### Summary of Updates

There has been a general revision to improve presentation and clarity, including changes to the structure and a new methods section.

