## Supplementary Materials for "THE CEREBELLUM CONVERTS INPUT DATA INTO A HYPER LOW-RESOLUTION GRANULE CELL CODE WITH SPATIAL DIMENSIONS: A HYPOTHESIS"

**Test of unreported ratios**

We explored the effect of varying model parameters for which physiological data are unreported. We did that to verify that the results are not dependent on hypothesis-favourable estimates.

The results are shown below. In **A**, the varied parameter is the number of mossy fibres with convergent input to a single Golgi cell dendrite, and in **B**, it is the number of Golgi cells with convergent input to a glomerulus. Both display the output of the conversion, representing the mean of Golgi cell rates received by each glomerulus in the field that receives ensemble output (so 700 values, the same as Figure 2, column 4). The ratios are shown above each panel. In other figures, ratios are 4:1 (mossy fibres that contact a Golgi cell basal dendrite) and 8–12:1 (Golgi cells that converge on a glomerulus).

We found that increasing either ratio condenses the plotted data range (representing tighter alignment of inhibition within a field), but there is a diminishing return. There is a stronger effect of increasing a low ratio and a progressively weakening effect at higher ratios. The only condition that returned a large range was a very low Golgi cell to glomerulus ratio. So: varying unknown parameters did not reveal a reason to discard or revise the model, provided that the physiological Golgi cell to glomerulus convergence ratio is not unexpectedly low.

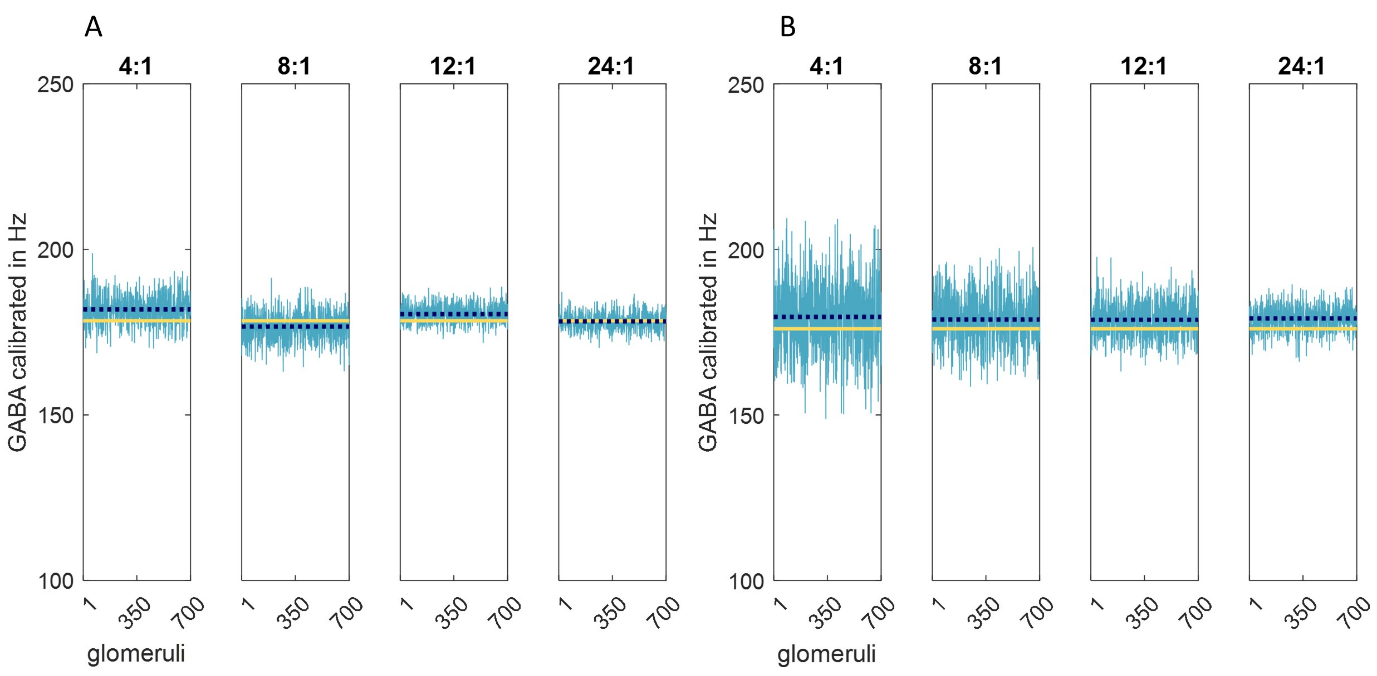

**Legend. Testing unreported parameters shows that reliability and precision are chiefly impacted at different steps in the computation.** All data were obtained in the same way as Fig 2 column 4, but we tested the effect of varying ratios for which physiological parameters have not been reported. Graphed data are accordingly the means of Golgi cell input rates to each of 700 glomeruli, the population of the middle field of a Golgi cell ensemble territory. Broken dark blue line: the mean of the means. The sampled distribution was generated in the same way as Fig 2 row 4. (The sampled distribution is the population of mossy fibre input rates to a row of 100 fields which includes the ensemble territory.) Yellow line: the mean of the sampled distribution. Varied ratios are the convergence ratio of mossy fibres onto a single Golgi cell basal dendrite (4:1 in Fig 2) and the convergence ratio of Golgi cells onto a single glomerulus is (a random number in the range 8–12:1 in Fig 2). In **A** we varied the first and in **B** the second. The ratios are shown above each panel. y axes cover a shorter range than Fig 2. A narrower range represents tighter within-field synchrony of inhibition received by granule cells. Findings were: (1) In both cases, there is a diminishing return of increasing the ratio. There is a modest return of higher ratios than Fig 2, selected to be physiologically plausible. (2) The blue mean varies in a small range around the yellow mean, but within that, varies at random each time the simulation is run. In (A), it varies from panel to panel because mossy fibre input rates to the simulated ensemble territory are resampled each time. In (B), that step in the computation is performed only once and the output used for all panels. Here, the blue mean is the same in all panels, but moves randomly when the simulation is rerun, in the same range as (A). So, reliability (reproducibility of the glomerular field mean) is impacted at the step of sampling of mossy fibre rates by Golgi cells, but not sampling of Golgi cell rates by glomeruli. Precision (narrowness of the glomerular range) is impacted by both ratios, but chiefly at the second step.
